# Host engineering for improved valerolactam production in *Pseudomonas putida*

**DOI:** 10.1101/660795

**Authors:** Mitchell G. Thompson, Luis E. Valencia, Jacquelyn M. Blake-Hedges, Pablo Cruz-Morales, Alexandria E. Velasquez, Allison N. Pearson, Lauren N. Sermeno, William A. Sharpless, Veronica T. Benites, Yan Chen, Edward E. K. Baidoo, Christopher J. Petzold, Adam M. Deutschbauer, Jay D. Keasling

## Abstract

*Pseudomonas putida* is a promising bacterial chassis for metabolic engineering given its ability to metabolize a wide array of carbon sources, especially aromatic compounds derived from lignin. However, this omnivorous metabolism can also be a hindrance when it can naturally metabolize products produced from engineered pathways. Herein we show that *P. putida* is able to use valerolactam as a sole carbon source, as well as degrade caprolactam. Lactams represent important nylon precursors, and are produced in quantities exceeding one million tons per year[1]. To better understand this metabolism we use a combination of Random Barcode Transposon Sequencing (RB-TnSeq) and shotgun proteomics to identify the *oplBA* locus as the likely responsible amide hydrolase that initiates valerolactam catabolism. Deletion of the *oplBA* genes prevented *P. putida* from growing on valerolactam, prevented the degradation of valerolactam in rich media, and dramatically reduced caprolactam degradation under the same conditions. Deletion of *oplBA*, as well as pathways that compete for precursors L-lysine or 5-aminovalerate, increased the titer of valerolactam from undetectable after 48 hours of production to ~90 mg/L. This work may serve as a template to rapidly eliminate undesirable metabolism in non-model hosts in future metabolic engineering efforts.

## 1 INTRODUCTION

*Pseudomonas putida* has attracted great attention as a potential chassis organism for metabolic engineering due in large part to its ability to metabolize a wide variety of carbon sources, particularly aromatic compounds that can be derived from lignin [2,3]. To more fully realize this vision, much effort has been put forth recently to better characterize the central metabolism of *P. putida* with updated genome-scale models [4], C^13^ flux experiments [5,6], and high-throughput fitness assays, which have all contributed to a more complete understanding of the bacterium [7,8]. Despite these advances, the metabolic capabilities of *P. putida* are not yet fully understood.

One consequence of this omnivorous metabolism is that *P. putida* possesses the ability to degrade or fully catabolize chemicals metabolic engineers seek to produce in the host. An ongoing challenge for *P. putida* host engineering will be to rapidly identify catabolic pathways of important target molecules and eliminate them from the genome. The recent report of a novel pathway for levulinic acid catabolism in *P. putida* KT2440 underscores the catabolic flexibility of the host, and an additional obstacle towards producing high product titer [8]. While challenging, this is not surprising; as a genus, *Pseudomonads* are well known for their ability to degrade a wide range of naturally occurring or xenobiotic chemicals [9,10].

Caprolactam and valerolactam are both important commodity chemicals used in the synthesis of nylon polymers, with global production of caprolactam reaching over four million metric tons [1]. Multiple efforts have been made to produce these chemicals biologically, with the titers of valerolactam approaching 1g/L in *Escherichia coli [1,11]*. The engineered pathway to valerolactam in *E. coli* converts L-lysine to 5-aminovalerate (5AVA) via DavBA, two genes endogenous to *P. putida*, and then cyclizes it via a promiscuous coA-ligase [1]. While the endogenous L-lysine catabolism catabolism of *P. putida* has been leveraged to produce the C5 diacid glutarate, there has yet to be any attempt to produce valerolactam in the bacterium [12]. A recent report that *Pseudomonas jessenii* can catabolize caprolactam suggested that catabolism of lactams could dramatically impact titers of valerolactam in *P. putida [13]*.

In this work we demonstrate that *P. putida* can utilize valerolactam as a sole carbon source, as well as degrade caprolactam. Using a combination of Random Barcode Transposon Sequencing (RB-TnSeq) and shotgun proteomics we were able to identify that OplBA, a predicted oxoprolinase, is responsible for this hydrolysis. By knocking out *oplBA* in addition to two pathways that compete for precursors we were able to dramatically increase the titers of valerolactam in *P. putida*.

## 2 RESULTS

### 2.1 Identification of a lactam hydrolase in *P. putida*

The hydrolysis product of valerolactam is 5AVA, an intermediate in L-lysine metabolism (Figure 1A). Growth curves of *P. putida* on valerolactam as a sole carbon source demonstrated that the bacterium was readily able to catabolize valerolactam and produce biomass, with growth similar to that on either 5AVA and glucose (Figure 1B). Initially, we attempted to identify the enzyme responsible for valerolactam hydrolysis using RB-TnSeq, a technique that has previously been used to identify novel enzymes in D-lysine metabolism [7]. RB-TnSeq measures the relative fitness of transposon mutant pools to infer gene function through changes in relative abundance of all non-essential genes in a bacterium under a selective condition [14,15]. Mutant pools of *P. putida* were grown on either 5AVA or valerolactam as a sole carbon source in an attempt to identify enzymes solely essential for growth on valerolactam. Results of RB-TnSeq experiments suggested that valerolactam was indeed being catabolized through the same pathway as 5AVA with both conditions showing significant defects in the *davTD* and *csiD-lghO* operons, the known catabolic route of 5AVA to the TCA cycle (Figure S1A). However, there were no genes that showed obvious fitness defects only under the valerolactam growth condition (Figure 1C).

**Figure 1:**
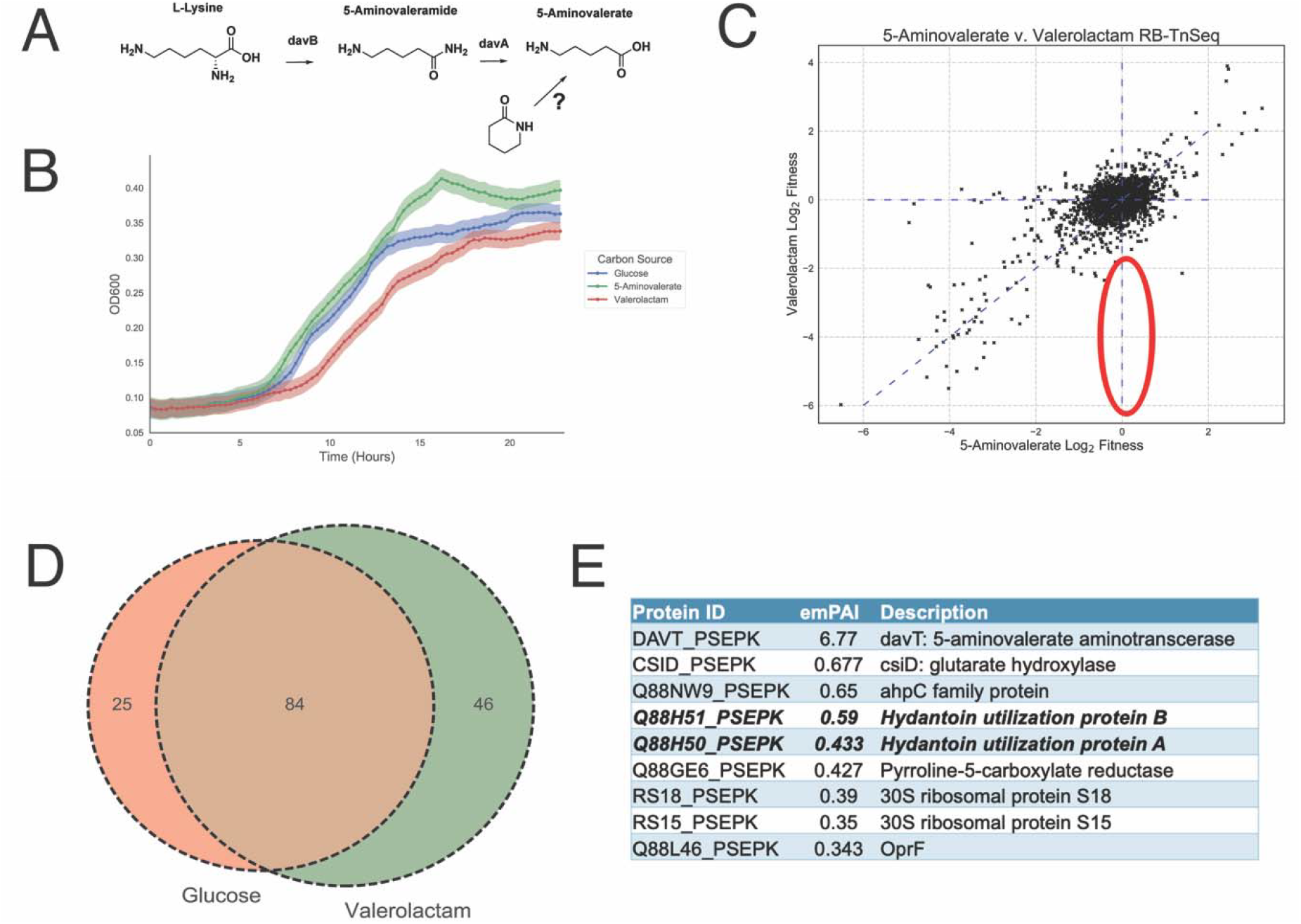
Identification of the *P. putida* valerolactam hydrolase: (A) Route of valerolactam catabolism through the L-lysine catabolic route of *P. putida* (B) Growth of *P. putida* in minimal medium supplemented with either 10 mM glucose, 5AVA, or valerolactam. Shaded area represents the 95% confidence interval (cI), n=3. (C) RB-TnSeq analysis of genome fitness assays of *P. putida* libraries grown on either 5AVA or valerolactam as a sole carbon source. Red oval shows the predicted fitness result of a valerolactam hydrolase. (D) Results of shotgun proteomics of proteins found in the supernatant of *P. putida* grown on either 10 mM glucose or 10 mM valerolactam as a sole carbon source. Venn diagram shows the number of proteins with an exponentially modified protein abundance index (emPAI) relative abundance above 0.1 shared or unique to each carbon source (E) Table shows the most abundant proteins specific to grown on valerolactam. OplA (Q88H50_PSEPK) and OplB (Q88H51_PSEPK) are in bold.

The inability of the RB-TnSeq approach to identify the hydrolytic enzyme could result from the enzyme being secreted from the cell. In this case, the secreted enzymes from cells containing the intact hydrolase gene produce 5AVA which can freely diffuse into hydrolase mutant cells, eliminating any selective pressure for lactam-based growth. To test whether the enzymes responsible for lactam hydrolysis may be secreted, cultures of *P. putida* were either grown on glucose or valerolactam as a sole carbon source, their supernatants filtered, concentrated, and then subjected to shotgun proteomics. Of the most abundant proteins in the supernatant, there were 25 proteins that were specific to glucose, while 46 were specific to valerolactam, with 86 proteins being shared between the two conditions (Figure 1D). Within the top five most abundant proteins in the valerolactam supernatant were OplB (Q88H51_PSEPK) and OplA (Q88H51_PSEPK), which together are annotated as the two subunits of a 5-oxoprolinase, orthologs of which have been suggested to participate in the caprolactam catabolism of *P. jessenii* (Figure 1E) [13]. Additional shotgun proteomics of cell pellets grown on either glucose, 5AVA, or valerolactam showed that OplB and OplA were more highly expressed in the presence of the lactam in comparison to the other carbon sources (Figure S1C). Interestingly, fitness data from two valerolactam RB-TnSeq experiments in *P. putida* KT2440 show *oplBA* mutants having no significant fitness defects (Figure S1B). Orthologs of *oplBA* are widely distributed across many bacteria including other attractive metabolic engineering chassis such as *Rhodococcus opacus*, and are nearly always located adjacent to one another on the genome (Figure S2).

### 2.2 Deletion of *oplBA* mitigates consumption of valerolactam and caprolactam

To confirm the role of OplBA in the catabolism of valerolactam, deletions were constructed of the *oplBA* locus in *P. putida* via homologous recombination. Growth of the mutant was compared to the wild type as well as to a deletion mutant of *davT*, which catalyzes the first step in 5AVA catabolism. All strains showed identical growth on glucose as a sole carbon source (Figure 2A). In the presence of 5AVA the *oplBA* deletion strain showed no growth defect compared to the wild-type, while the *davT* mutant predictably was unable to grow (Figure 2B). However, on valerolactam only the wild type strain grew, while both the *oplBA* and *davT* mutants showed no measurable growth after 40 hours (Figure 2C). These results suggest that OplBA is the sole enzyme responsible for the conversion of valerolactam to 5AVA under these conditions.

**Figure 2:**
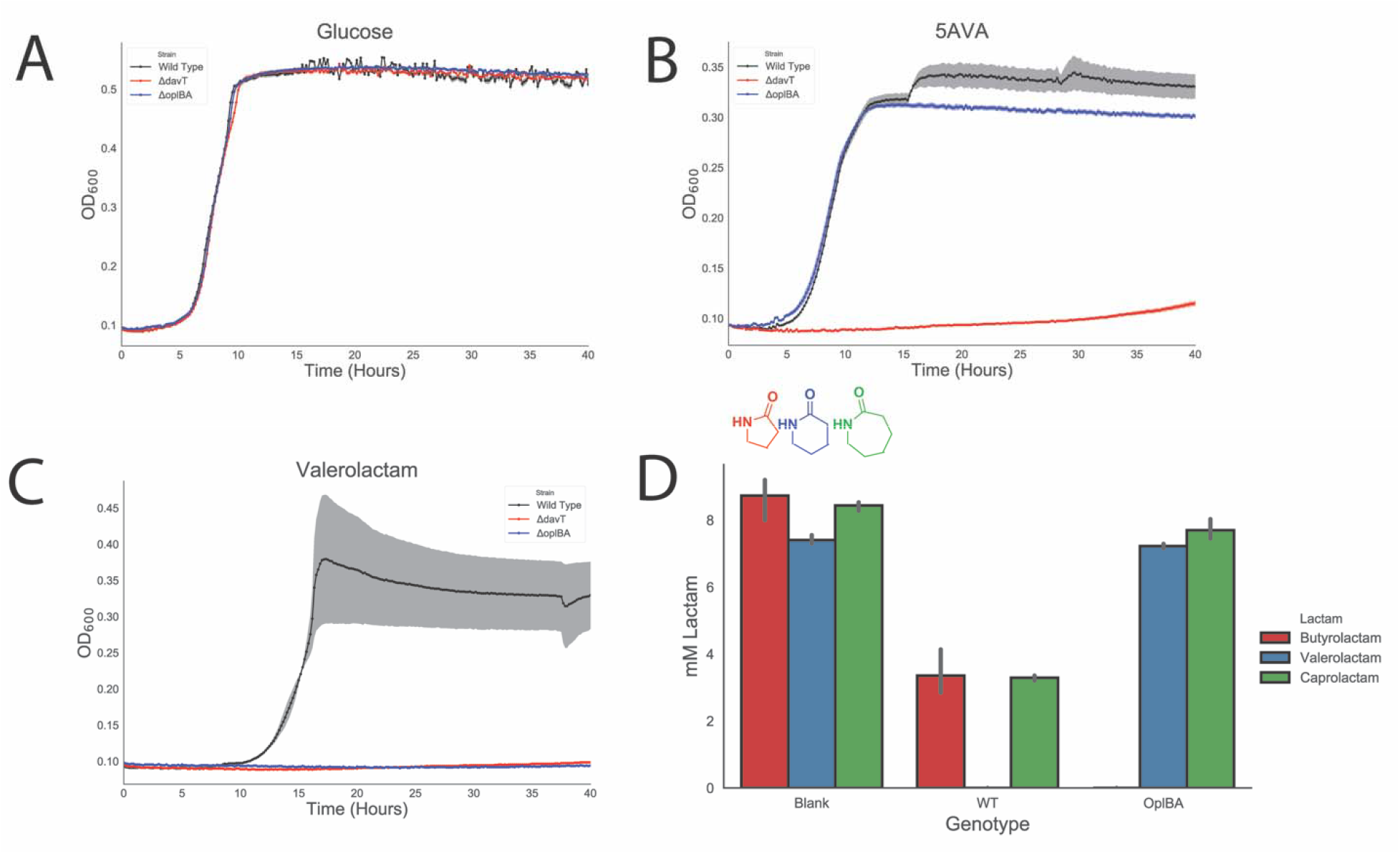
OplBA controls valerolactam and caprolactam degradation in *P. putida*. Growth of wild-type, Δ*davT*, or Δ*oplBA* in minimal media supplemented with either 10 mM glucose (A), 5-aminovaleroate (B), or valerolactam (C). (D) Remaining butyrolactam, valerolactam or caprolactam in LB media after 24-hour incubation with no *P. putida*, wild-type, or a Δ*oplBA* mutant.

While *P. putida* KT2440 can utilize valerolactam as a sole carbon source, it is unable to grow on caprolactam (data not shown). In order to determine whether the OplBA of *P. putida* is capable of degrading other lactams, wild type and Δ*oplBA* strains were grown in LB medium supplemented with 10 mM caprolactam, valerolactam, or butyrolactam for 24 hours after which the remaining lactam concentration was compared to an uninoculated medium control. Wild type *P. putida* consumed all detectable valerolactam within 24 hours, and less than 50% of both butyrolactam and caprolactam remained compared to the uninoculated control (Figure 2D). There was no significant decrease in amount of valerolactam in the Δ*oplBA* cultures compared to the uninoculated control (t-test of *p*=0.178), though there was a slight but significant decrease of 0.74 mM caprolactam (t-test of *p*=0.033). No butyrolactam remained in the Δ*oplBA* culture after 24 hours (Figure 2D). This result was surprising as the annotated substrate of OplBA, 5-oxoproline, has the same ring size as butyrolactam. Previous work has shown that homologs of OplBA are ATP-dependent amidohydrolases that hydrolyze 5-oxoproline [16]. However, when purified OplBA was incubated with valerolactam in addition to ATP and magnesium we observed no hydrolysis relative to boiled enzyme controls (Figure S3).

### 2.3 Host engineering for increased valerolactam production

The published pathway for the production of valerolactam from L-lysine in *E. coli* utilized the formation of 5AVA via the *davBA* pathway native to *P. putida*, followed by cyclization to the lactam via a promiscuous acyl-coA ligase from *Streptomyces aizunensis* (Figure 4A) [1]. To produce valerolactam in *P. putida*, not only will the production pathway need to be overexpressed, but the *oplBA* locus and pathways that compete for 5AVA must be eliminated. Loss of flux to valerolactam occurs through two known competing pathways: the Alr-mediated isomerization to D-lysine or catabolism of 5AVA to glutarate via the action of DavTD (Figure 3A) [7].

**Figure 3:**
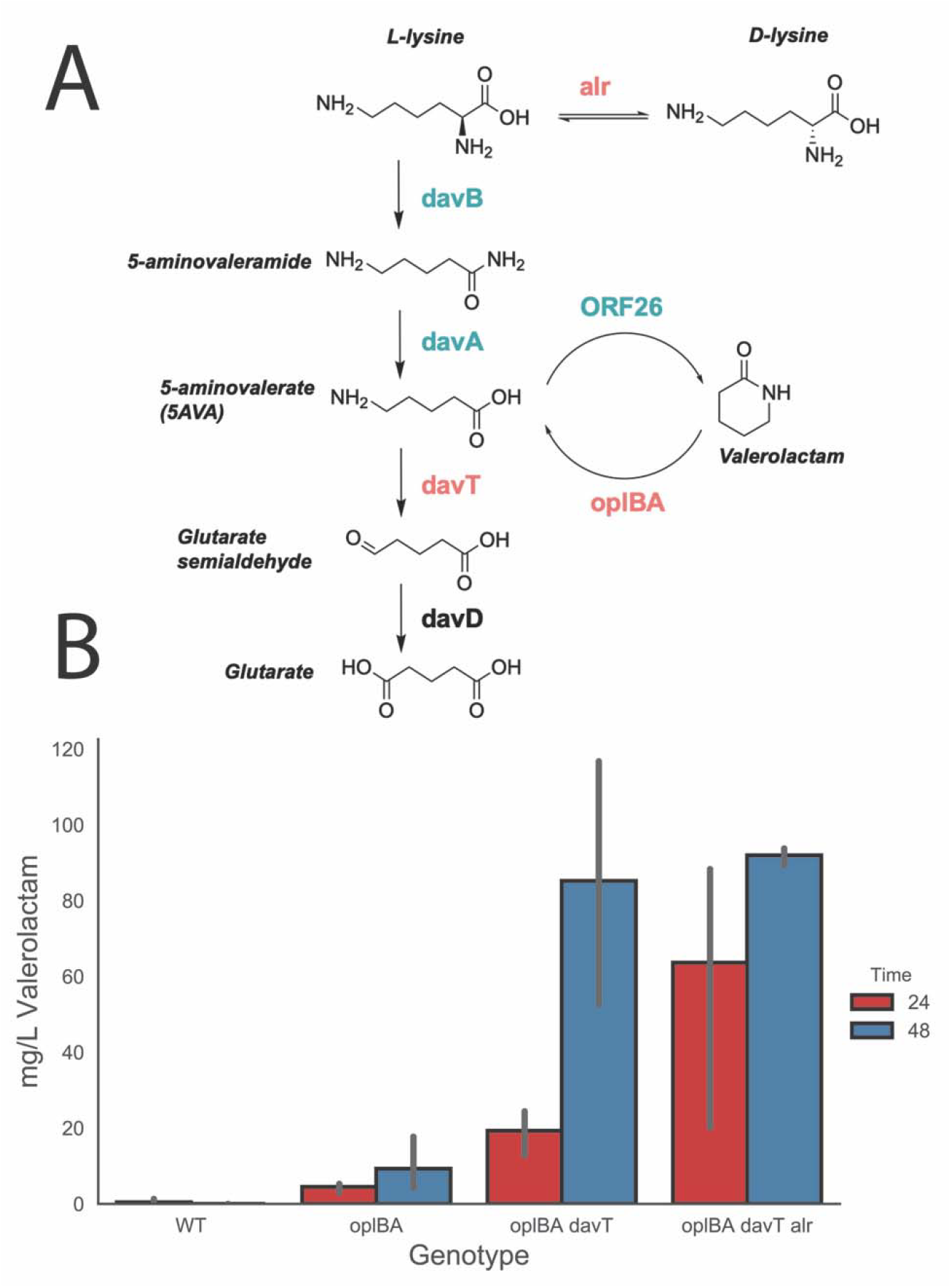
Production of valerolactam in *P. putida*: (A) Design of valerolactam overproducing *P. putida*. The biosynthetic pathway genes are shown in green and were overexpressed heterologously from a pBBR ori plasmid using an arabinose inducible promoter. Pathways that catabolize products or divert precursors are shown in red. (B) Valerolactam production from different *P. putida* strains grown in LB medium supplemented with 25 mM L-lysine and 0.2% (w/v) arabinose at 24 and 48 hours post inoculation. Error bars show 95% cI, n=3.

To investigate the relative contributions of pathways that contribute to either lactam hydrolysis or loss of substrate, we expressed the *davBA-ORF26* pathway via a arabinose-inducible broad host range vector pBADT in backgrounds where these pathways had been sequentially deleted. Wild type *P. putida* produced 0.43 mg/L valerolactam after 24 hours, but no valerolactam could be detected after 48 hours, presumably due to host consumption (Figure 3B). Simple deletion of the *oplBA* locus resulted in a 10-fold increase of production at 24 hours to 4.47 mg/L and a 48-hour titer of 9.27 mg/L (Figure 3B). Additional deletion of *davT* resulted in an increase of titer to 19.29 mg/L and 85.19 mg/L, and by deleting the amino acid racemase *alr* titers increased to 63.66 mg/L and 91.97 mg/L at 24 and 48 hours, respectively (Figure 3B).

## 3 DISCUSSION

Recent economic analyses highlight the necessity of lignin valorization to create a sustainable bioeconomy [17,18]. With its robust aromatic metabolism combined with novel methods of biomass deconstruction, *P. putid*a has great potential to convert lignocellulosic biomass to value-added products [19,20]. Though the ability of *P. putida* to catabolize many carbon sources is often viewed as an asset, it has well documented metabolic pathways to degrade many compounds that metabolic engineers may wish to produce such as levulinic acid [8], various alcohols [21], and the diacid glutarate [12]. As these catabolic phenotypes are encountered it will be critical to rapidly identify the offending genomic loci.

The recent surge in development of functional genomics techniques has dramatically increased the throughput at which we can identify the genetic basis of unknown metabolism. RB-TnSeq has been used to uncover novel glutarate and levulinic acid metabolism, though was ineffective at identifying the *oplBA* locus [7,8]. Proteomics techniques have also grown more robust, and were used in *P. jessenii* to predict a route of caprolactam catabolism [13,22]. Here, proteomics also proved to be an effective means of identifying the enzyme responsible for the hydrolysis of the lactam. Our proteomics results showed that OplBA was specifically expressed when grown on valerolactam, but not 5AVA. These results suggest that *P. putida* may have lactam-specific transcription factors which could be developed into valuable tools for metabolic engineering if identified.

The inability of RB-TnSeq to identify these genes is curious as single deletion mutants were unable to grow on valerolactam. A possible explanation for this is that OplBA may be secreted, which would create a public pool of 5AVA which *oplBA* mutants could still utilize. However, OplBA orthologs have been shown to be ATP-dependent [16], which would be inconsistent with this extracellular localization. Unfortunately our attempts to characterize OplBA *in vitro* were unsuccessful, preventing us from identifying the substrate requirements of the enzyme. Though we were unable to reconstitute the activity of OplBA *in vitro*, deletion of *oplBA* did not prevent the degradation of butyrolactam which is a 5-membered lactam ring. The annotated function of the *oplBA* loci is a 5-oxoprolinase, which hydrolyzes the 5-membered lactam ring of 5-oxoproline. These results may suggest that the OplBA may function naturally as something other than a 5-oxoprolinase. More work will be necessary to resolve the results of our RB-TnSeq and feeding experiments to elucidate the biochemical requirements of OplBA as well as to better understand its cellular localization.

Without deleting *oplBA P. putida* is able to metabolize up to 10 mM valerolactam in rich media after 24 hours, and simple deletion of these genes increases production 10-fold after 24 hours of fermentation. Subsequent deletion of the 5AVA transaminase *davT* and the racemase *alr* resulted in 48 hour titers of ~90 mg/L, whereas there was no detectable valerolactam production in wild-type cultures at this time point. To achieve these titers we fed in 25 mM L-lysine (3.65 g/L). Previous work in *E. coli* achieved titers of ~200 mg/L by feeding 1 g/L lysine and ~300 mg/L by feeding 5 g/L after 48 hours [1]. Our results indicate that with significant host engineering, *P. putida* can produce titers approaching those of model organisms. Optimization of pathway expression could narrow this gap even further and should be a focus of future efforts.

While this initial work is encouraging, it still requires the feeding of L-lysine in rich media for conversion to valerolactam. Ideally, engineering *P. putida* would be able to metabolize lignin hydrolysis products directly to L-lysine on the way to the final product. A great deal of work has been conducted to elucidate the complex catabolism of lysine in *P. putida [7,12]*, yet relatively little work has been done to increase flux to lysine within the bacterium. While there has been little to divert flux to L-lysine in *P. putida*, there is a wealth of evidence in other bacteria where high titers of intracellular lysine have been achieved [23,24]. The ever increasing body of research to characterize the sprawling metabolism of *P. putida* will greatly aid in future efforts of efficient production of valerolactam from lignocellulosic feedstocks.

## 4 METHODS

### 4.1 Media, chemicals, and culture conditions

General *E. coli* cultures were grown in Luria-Bertani (LB) Miller medium (BD Biosciences, USA) at 37 °C while *P. putida* was grown at 30 °C. When indicated, *P. putida* and *E. coli* were grown on modified MOPS minimal medium [25]. Cultures were supplemented with kanamycin (50□mg/L, Sigma Aldrich, USA), gentamicin (30□mg/L, Fisher Scientific, USA), or carbenicillin (100mg/L, Sigma Aldrich, USA), when indicated. All other compounds were purchased through Sigma Aldrich (Sigma Aldrich, USA).

### 4.2 Strains and plasmids

All bacterial strains and plasmids used in this work are listed in Table 1. All strains and plasmids created in this work are available through the public instance of the JBEI registry. (https://public-registry.jbei.org/folders/456). All plasmids were designed using Device Editor and Vector Editor software, while all primers used for the construction of plasmids were designed using j5 software [26–28]. Plasmids were assembled via Gibson Assembly using standard protocols [29], or Golden Gate Assembly using standard protocols [30]. Plasmids were routinely isolated using the Qiaprep Spin Miniprep kit (Qiagen, USA), and all primers were purchased from Integrated DNA Technologies (IDT, Coralville, IA).

**Table 1:** Strains and plasmids used in this study

| Strain | JBEI Part ID | Reference |
| --- | --- | --- |
| E. coli DH10B |  | [31] |
| E. coli BL21(DE3) |  | Novagen |
| P. putida KT2440 |  | ATCC 47054 |
| *P. putida* Δ*davT* | | [32] |
| P. putida Δ*oplBA* | JBEI-104285 | This work |
| P. putida Δ*oplBA*Δ*davT* | JBEI-104286 | This work |
| P. putida Δ*oplBA*Δ*davT*Δ*alr* | JBEI-104287 | This work |
| **Plasmids** |  |  |
| pET28 |  | Novagen |
| pET28 *oplA* | JBEI-104336 | This work |
| pET28 *oplB* | JBEI-104337 | This work |
| pBADT |  | [33] |
| pBADT-*davBA*-ORF26 | JBEI-104356 | This work |
| pMQ30 |  | [34] |
| pMQ30 *oplBA* | JBEI-104355 | This work |
| pMQ30 *alr* | JBEI-104354 | This work |
| pMQ30 *davT* |  | [32] |

### 4.3 Plate based growth assays

Growth studies of bacterial strains were conducted a microplate reader kinetic assays. Overnight cultures were inoculated into 10 mL of LB medium from single colonies, and grown at 30□°C. These cultures were then washed twice with MOPS minimal media without any added carbon and diluted 1:100 into 500 μL of MOPS medium with 10 mM of a carbon source in 48-well plates (Falcon, 353072). Plates were sealed with a gas-permeable microplate adhesive film (VWR, USA), and then optical density and fluorescence were monitored for 48□hours in an Biotek Synergy 4 plate reader (BioTek, USA) at 30□°C with fast continuous shaking. Optical density was measured at 600□nm.

### 4.4 Production Assays and Lactam Quantification

To assess valerolactam production in strains of *P. putida* overnight cultures of strains harboring pBADT-*davBA-ORF26* were grown in 3 mL of LB supplemented with kanamycin and grown at 30L□°C. Production cultures of 10 mL of LB supplemented with kanamycin, 25mM L-lysine, and 0.2% w/v arabinose were then inoculated 1:100 with overnight cultures and then grown at 30□°C shaking at 250 rpm. Samples for valerolactam production were taken at 24 and 48 hours post-inoculation, with 200 μL of culture being quenched with an equal volume of ice cold methanol and then stored at −20 °C until analysis.

For measurement of lactams, liquid chromatographic separation was conducted at 20°C with a Kinetex HILIC column (50-mm length, 4.6-mm internal diameter, 2.6-µm particle size; Phenomenex, Torrance, CA) using a 1260 Series HPLC system (Agilent Technologies, Santa Clara, CA, USA). The injection volume for each measurement was 5 µL. The mobile phase was composed of 10 mM ammonium formate and 0.07% formic acid in water (solvent A) and 10 mM ammonium formate and 0.07% formic acid in 90% acetonitrile and 10% water (solvent B) (HPLC grade, Honeywell Burdick & Jackson, CA, USA). High purity ammonium formate and formic acid (98-100% chemical purity) were purchased from Sigma-Aldrich, St. Louis, MO, USA. Lactams were separated with the following gradient: decreased from 90%B to 70%B in 2 min, held at 70%B for 0.75 min, decreased from 70%B to 40%B in 0.25 min, held at 40%B for 1.25 min, increased from 40%B to 90%B for 0.25 min, held at 90%B for 1 min. The flow rate was varied as follows: 0.6 mL/min for 3.25 min, increased from 0.6 mL/min to 1 mL/min in 0.25 min, and held at 1 mL/min for 2 min. The total run time was 5.5 min.

The HPLC system was coupled to an Agilent Technologies 6520 quadrupole time-of-flight mass spectrometer (QTOF MS) with a 1:6 post-column split. Nitrogen gas was used as both the nebulizing and drying gas to facilitate the production of gas-phase ions. The drying and nebulizing gases were set to 12 L/min and 30 lb/in^2^, respectively, and a drying gas temperature of 350°C was used throughout. Fragmentor, skimmer and OCT 1 RF voltages were set to 100 V, 50 V and 300 V, respectively. Electrospray ionization (ESI) was conducted in the positive-ion mode for the detection of [M + H]^+^ ions with a capillary voltage of 4000 V. The collision energy voltage was set to 0 V. MS experiments were carried out in the full-scan mode (75–1100 *m/z*) at 0.86 spectra/s. The QTOF-MS system was tuned with the Agilent ESI-L Low concentration tuning mix in the range of 50-1700 *m/z*. Lactams were quantified by comparison with 8-point calibration curves of authentic chemical standards from 0.78125 µM to 100 µM. R^2^ coefficients of ≥0.99_[EB1]_ were achieved for the calibration curves. Data acquisition was performed by Agilent MassHunter Workstation (version B.05.00), qualitative assessment by Agilent MassHunter Qualitative Analysis (version B.05.00 or B.06.00), and data curation by Agilent Profinder (version B.08.00)

### 4.5 RB-TnSeq and Proteomics Analysis

RB-TnSeq experiments utilized *P. putida* library JBEI-1 which has been described previously [7]. Libraries of JBEI-1 were thawed on ice, diluted into 25 mL of LB medium with kanamycin and then grown to an OD_600_ of 0.5 at 30□°C at which point three 1 mL aliquots were removed, pelleted, and stored at −80□°C. Libraries were then washed once in MOPS minimal medium with no carbon source, and then diluted 1:50 in MOPS minimal medium with 10 mM valerolactam. Cells were grown in 500 μL of medium in 48-well plates (Falcon, 353072). Plates were sealed with a gas-permeable microplate adhesive film (VWR, USA), and then grown at 30□°C in a Tecan Infinite F200 microplate reader (Tecan Life Sciences, San Jose, CA), with shaking at 200 rpm. Two 500 μL aliquots were combined, pelleted, and stored at −80□°C until BarSeq analysis, which was performed as previously described [8,14]. All fitness data is publically available at http://fit.genomics.lbl.gov.

Secreotomes of *P. putida* were prepared by growing 500 mL of culture in MOPS minimal medium supplemented with either 10 mM glucose or 10 mM valerolactam for 24 hours at 30□°C, which were subsequently pelleted and filtered through a 0.22 μm filter and then concentrated 100x via a 10 kD MW cutoff filter. Cultures for intracellular proteomics analysis were grown in 10 mL cultures in the same conditions on either glucose, 5AVA, or valerolactam and were then pelleted and stored at −80□°C until sample workup and proteomic analysis. Proteins from secreted and intracellular samples were desalted and isolated using a variation of a previously-described chloroform/methanol extraction protocol [35]. For secreted proteins, 100-200 μL of the concentrated protein sample was used; for intracellular samples, cell pellets were thoroughly resuspended in 100 μL HPLC water. Then, the following reagents were added to each sample in sequential order with thorough vortexing after each addition: 400 μL of HPLC grade methanol, 100 μL of HPLC grade chloroform, 300 μL of HPLC grade water. Samples were centrifuged for 1 minute at ~21,000g in order to promote phase separation. After centrifugation, the entirety of the top layer (water and methanol) was removed and discarded, leaving on the protein pellet and chloroform layer remaining. Another 300 μL of HPLC grade methanol was added, then the samples were vortexed and centrifuged again for 2 minutes at ~21,000g. The remaining liquid was then removed and discarded, and the cell pellets were allowed to dry in a fume hood for 5 minutes. Protein pellets were then resuspended in freshly-prepared 100mM ammonium bicarbonate buffer in HPLC water containing 20% HPLC methanol. Protein concentrations in the resuspended samples were quantified using a DC Protein Assay Kit (Bio-Rad Laboratories, Hercules, CA). After quantification, 100 μg of protein was transferred to a PCR strip and tris(2-carboxyethyl)phosphine was added to a final concentration of 5mM. Samples were incubated at 22 °C for 30 minutes, after which iodoacetamide was added (final concentration 10mM). Samples were again incubated at 22 °C in the dark for 30 minutes. Finally, trypsin was added to a final ratio of 1:25 w/w trypsin:sample, and samples were digested at 37 °C for 5-8 hours before being transferred to conical LC vials for LC-MS analysis. Peptides prepared for shotgun proteomic experiments were analyzed by using an Agilent 6550 iFunnel Q-TOF mass spectrometer (Agilent Technologies, Santa Clara, CA) coupled to an Agilent 1290 UHPLC system as described previously [36]. 20 μg of peptides were separated on a Sigma–Aldrich Ascentis Peptides ES-C18 column (2.1 mm × 100 mm, 2.7 μm particle size, operated at 60 °C) at a 0.400 mL/min flow rate and eluted with the following gradient: initial condition was 98% solvent A (0.1% formic acid) and 2% solvent B (99.9% acetonitrile, 0.1% formic acid). Solvent B was increased to 35% over 30 min, and then increased to 80% over 2 min, then held for 6 min, followed by a ramp back down to 2% B over 1 min where it was held for 4 min to re-equilibrate the column to original conditions. Peptides were introduced to the mass spectrometer from the LC by using a Jet Stream source (Agilent Technologies) operating in positive-ion mode (3,500 V). Source parameters employed gas temp (250°C), drying gas (14 L/min), nebulizer (35 psig), sheath gas temp (250°C), sheath gas flow (11 L/min), VCap (3,500 V), fragmentor (180 V), OCT 1 RF Vpp (750 V). The data were acquired with Agilent MassHunter Workstation Software, LC/MS Data Acquisition B.06.01 operating in Auto MS/MS mode whereby the 20 most intense ions (charge states, 2–5) within 300–1,400 m/z mass range above a threshold of 1,500 counts were selected for MS/MS analysis. MS/MS spectra (100–1,700 m/z) were collected with the quadrupole set to “Medium” resolution and were acquired until 45,000 total counts were collected or for a maximum accumulation time of 333 ms. Former parent ions were excluded for 0.1 min following MS/MS acquisition. The acquired data were exported as mgf files and searched against the latest *P. putida* KT2440 protein database with Mascot search engine version 2.3.02 (Matrix Science). The resulting search results were filtered and analyzed by Scaffold v 4.3.0 (Proteome Software Inc.).

### 4.6 Protein Purification and Biochemical Analysis of OplBA

Both *oplB* and *oplA* were cloned into the expression vector pET28 harboring N-terminal 6x-histidine purification tags. Protein expression and purification were carried out as described previously [7]. To characterize activity of OplBA we used conditions that were previously described to characterize 5-oxoprolinase with minor changes [37]. Briefly, 10 μM of each OplB and OplA or boiled controls were incubated for 4 hours at 30 °C with 2 mM valerolactam in 100 mM Tris-HCl pH 7.0, 4 mM MgCl_2_, with or without 2 mM ATP. Reactions were quenched with ice-cold methanol, filtered through a 3kDa MWCO filter, diluted 40-fold with 50% methanol/50% H2O, and stored at −20 °C until analysis via LC-MS.

### 4.7 Bioinformatic Analyses

All statistical analyses were carried out using either the Python Scipy or Numpy libraries [38,39]. For the phylogenetic reconstructions, the best amino acid substitution model was selected using ModelFinder as implemented on IQ-tree [40] phylogenetic trees were constructed using IQ-tree, nodes were supported with 10,000 bootstrap replicates. The final tree figures were edited using FigTree v1.4.3 (http://tree.bio.ed.ac.uk/software/figtree/). Orthologous syntenic regions of OplBA were identified with CORASON-BGC [41] and manually colored and annotated.

## Acknowledgements

We thank Jesus Barajas for his careful reading of this manuscript and helpful suggestions during preparation. We also thank Morgan Price for assistance in analyzing RB-TnSeq data. We would like to thank the Amgen Scholars program for providing funding for Alexandria Velasquez.

This work was part of the DOE Joint BioEnergy Institute (https://www.jbei.org) supported by the U. S. Department of Energy, Office of Science, Office of Biological and Environmental Research, supported by the U.S. Department of Energy, Energy Efficiency and Renewable Energy, Bioenergy Technologies Office, through contract DE-AC02-05CH11231 between Lawrence Berkeley National Laboratory and the U.S. Department of Energy. The views and opinions of the authors expressed herein do not necessarily state or reflect those of the United States Government or any agency thereof. Neither the United States Government nor any agency thereof, nor any of their employees, makes any warranty, expressed or implied, or assumes any legal liability or responsibility for the accuracy, completeness, or usefulness of any information, apparatus, product, or process disclosed, or represents that its use would not infringe privately owned rights. The United States Government retains and the publisher, by accepting the article for publication, acknowledges that the United States Government retains a nonexclusive, paid-up, irrevocable, worldwide license to publish or reproduce the published form of this manuscript, or allow others to do so, for United States Government purposes. The Department of Energy will provide public access to these results of federally sponsored research in accordance with the DOE Public Access Plan (http://energy.gov/downloads/doe-public-access-plan)

## Contributions

Conceptualization, M.G.T.; Methodology, M.G.T., L.E.V.,J.M.B, P.C.M.,E.E.K.B.; Investigation, M.G.T., L.E.V., J.M.B, P.C.M, V.T.B, W.A.S., A.N.P., A.E.V.; Writing – Original Draft, M.G.T.; Writing – Review and Editing, All authors.; Resources and supervision, C.J.P., A.M.D., J.D.K.

---

**Figure S1:**
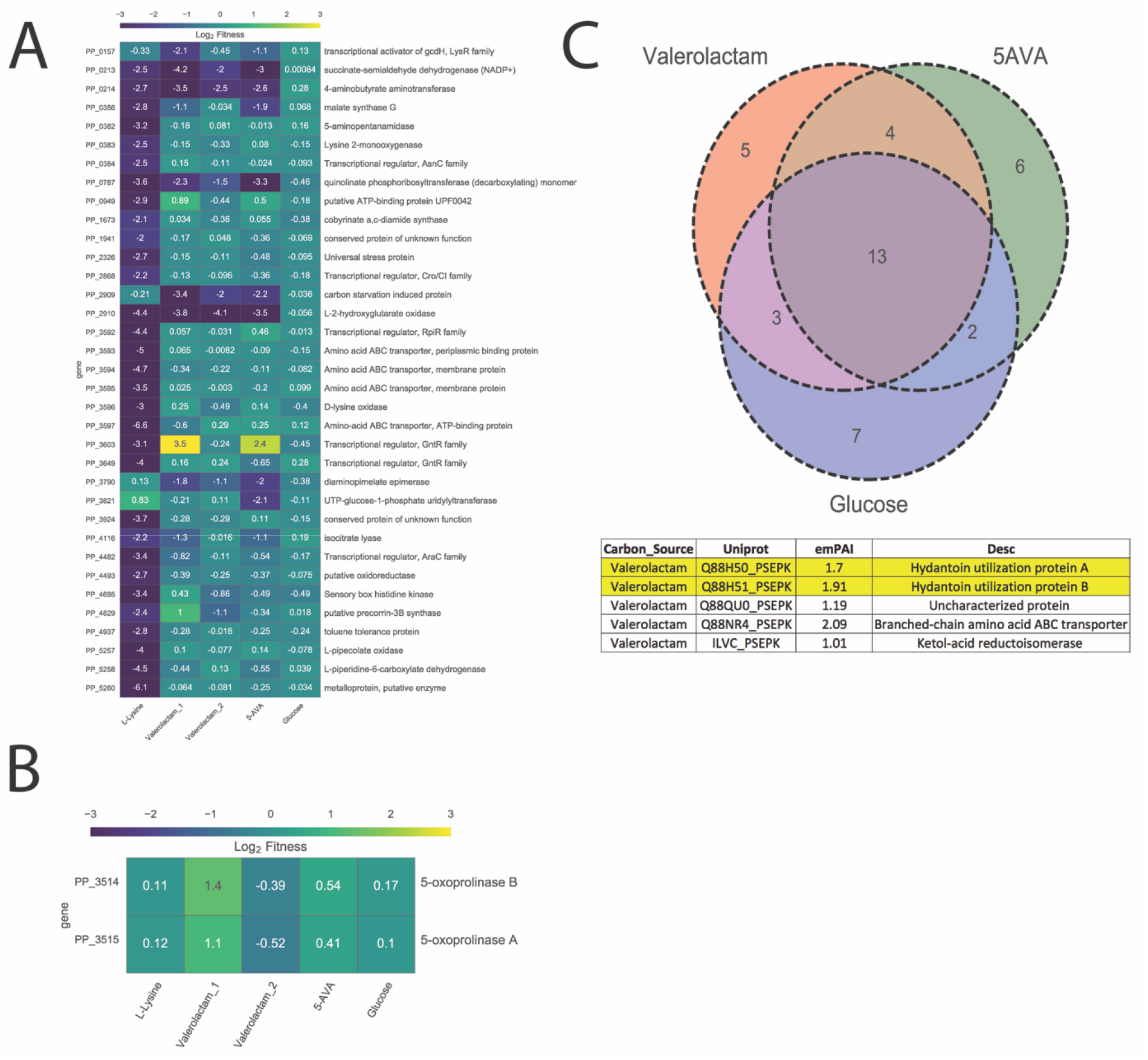
RB-TnSeq and Cellular Shotgun Proteomics Results. (A) Genes that show significant (t < −4) and large (fitness < −2) fitness defects specific to either L-lysine, 5AVA, or valerolactam, but not glucose. All non-valerolactam fitness experiments are from Thompson et. al 2019. (B) Fitness of the *oplA* and *oplB* genes on all carbon sources (C) Venn diagram showing the number of specific proteins found in the 100 most abundant proteins found within *P. putida* grown on either glucose, valerolactam, or 5-aminovalerate, based on emPAI. Below we can see the 5 most abundant proteins specific to growth on valerolactam. OplA and OplB are highlighted.

**Figure S2:**
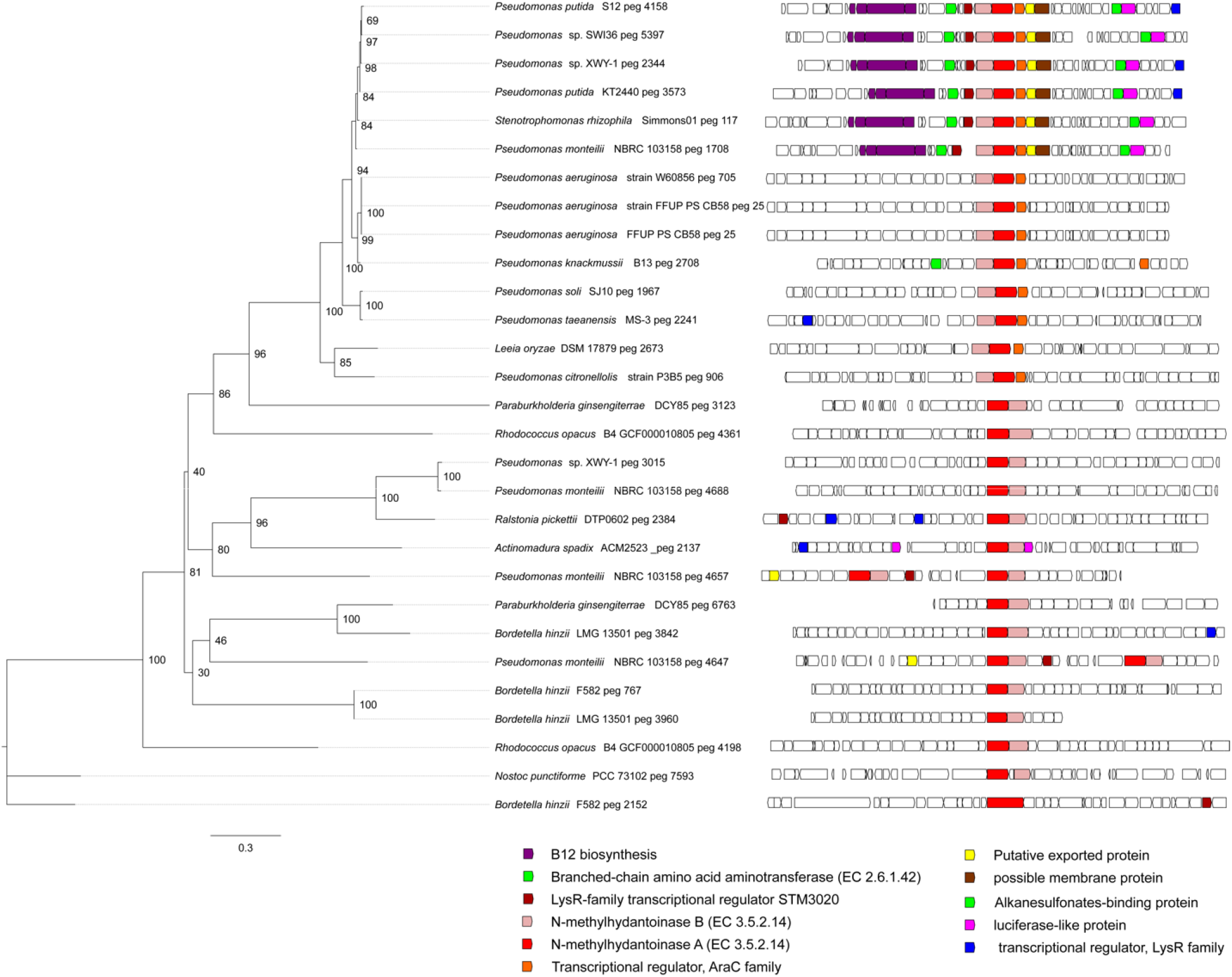
Distribution of OplBA orthologs: Phylogenomics of selected OplBA homologs across bacteria. The boxes represent the gene neighborhood for each homolog. The genes have been colored to represent their annotated functions.

**Figure S3:**
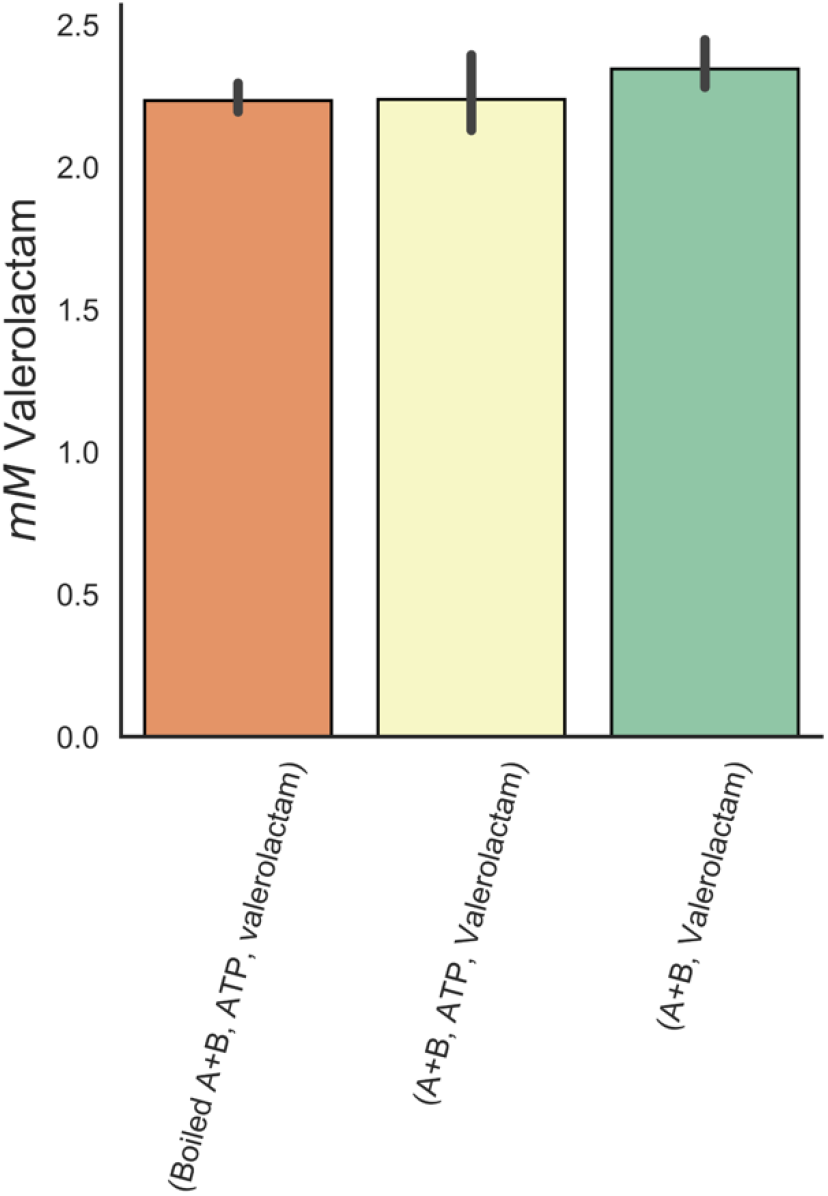
Biochemical characterization of OplAB. When purified OplA and OplB were incubated in the presence of valerolactam with or without ATP for 4 hours, there was no significant decrease in lactam concentration compared to boiled enzyme control. Error bars represent 95% cI, n=3.

